# Combining Unity with machine vision to create low latency, flexible, and simple virtual realities

**DOI:** 10.1101/2024.02.05.579029

**Authors:** Yuri Ogawa, Raymond Aoukar, Richard Leibbrandt, Jake S Manger, Zahra M Bagheri, Luke Turnbull, Chris Johnston, Pavan K Kaushik, Jan M Hemmi, Karin Nordström

## Abstract

1. In recent years, virtual reality arenas have become increasingly popular for quantifying visual behaviors. By using the actions of a constrained animal to control the visual scenery, the animal is provided the perception of moving through a simulated environment. As the animal is constrained in space, this allows detailed behavioral quantification. Additionally, as the world is generally computer-generated this allows for mechanistic quantification of visual triggers of behavior.
2. We created a novel virtual arena combining machine vision with the gaming engine Unity. For tethered flight, we enhanced an existing multi-modal virtual reality arena, MultiMoVR (Kaushik et al., 2020) but tracked hoverfly wing movements using DeepLabCut-live (DLC-live, Kane et al., 2020). For trackball experiments, we recorded the motion of a ball that a tethered crab was walking on using FicTrac (Moore et al., 2014). In both cases, real-time tracking was interfaced with Unity to control the movement of the tethered animals’ avatars in the virtual world. We developed a user-friendly Unity Editor interface, CAVE, to simplify experimental design and data storage without the need for coding.
3. We show that both the DLC-live-Unity and the FicTrac-Unity configurations close the feedback loop effectively with small delays, less than 50 ms. Our FicTrac-Unity integration highlighted the importance of closed-loop feedback by reducing behavioral artifacts exhibited by the crabs in open-loop scenarios. We show that *Eristalis tenax* hoverflies, using the DLC-live-Unity integration, navigate towards flowers. The effectiveness of our CAVE interface is shown by implementing experimental sequencing control based on avatar proximity to virtual structures.
4. Our results show that combining Unity with machine vision tools such as DLC-live and FicTrac provides an easy and flexible virtual reality (VR) environment that can be readily adjusted to new experiments and species. This can be implemented programmatically in Unity, or by using our new tool CAVE, which allows users to design and implement new experiments without programming in code. We provide resources for replicating experiments and our interface CAVE via GitHub, together with user manuals and instruction videos, for sharing with the wider scientific community.

## Introduction

Many animals use visual information to control fast and dynamic behaviors. Such behaviors include course stabilization and obstacle avoidance during navigation, homing, conspecific interactions, and predator avoidance. These behaviors are often studied in the laboratory, as this provides better control of the parameter space. However, a draw-back of such experiments is that as they are often so constrained that the ecological relevance comes into question (Gomez-Marin and Ghazanfar, 2019). Alternatively, visual behaviors can be studied under completely natural conditions in the field. While this produces unconstrained, naturalistic behaviors, the lack of experimental control makes mechanistic analyses difficult.

Due to recent technical advances, virtual reality (VR) arenas have become more popular for behavioral quantification. They are more likely to produce more naturalistic visual behaviors, while still being constrained enough to provide highly detailed behavioral quantifications. In VR, the actions of a constrained animal control changes of a simulated environment, thus providing the perception of moving within the simulated world (Dombeck and Reiser, 2012). As the animal is tethered, and thus not physically moving through space, monitoring the animal and quantifying its responses is greatly simplified. In addition, as the visual world is computer-generated, it is easy to manipulate, and thus allows systematic control over the parameter space, facilitating identification of visual triggers of behavior.

Insect VR include closed loop tethered flight arenas, which originally used a torque meter to quantify yaw turns (Dickinson et al., 1993). They now use the difference between the left and the right wing’s peak downstroke angle, or wing beat amplitude (WBA), which are strongly correlated with the yaw torque (Tammero et al., 2004), to control yaw steering motion (see e.g. Maimon et al., 2010). Alternative approaches include tracking the abdomen of moths with optical sensors, as the abdominal movements follow the forewing asymmetry (Gray et al., 2002). However, these arenas are often focused on yaw rotational motion, and the commonly used modular LED arenas (Reiser and Dickinson, 2008) often lack the 3D cues associated with translation (but see e.g. Ruiz and Theobald, 2020). This is important, as flying insects perform both rotational and translational behaviors (Geurten et al., 2012), such as forward motion and sideslip, which require thrust measurements.

Other VR arenas allow translational motion, including, for example, TrackFly (Fry et al., 2009) and FreemoVR (Stowers et al., 2017), developed for freely moving animals, and validated in zebrafish, flies, and mice, where the visual surround is updated based on the animal’s current position (see also Cruz et al., 2021; Pokusaeva et al., 2023). While these allow real-time closing of the loop based on the animal’s spatial position, it can be difficult to analyze finer behaviors offline. For example, many flying insects move the head relative to the body (Cellini et al., 2022; Talley et al., 2023), the antennas actively move (Sant and Sane, 2018), and the legs might perform landing movements (Shen and Sun, 2017). In this respect, the Antarium is an interesting VR framework, where a tethered ant’s walking on an air-supported ball controls both (yaw) rotational and translational motion through a Unity generated virtual environment (Kócsi et al., 2020). Analogously, a recently developed multi-sensory tethered flight arena, MultiMoVR, allows both yaw rotations and translations in the virtual environment (Kaushik et al., 2020).

For VR to provide an immersive experience it is important to close the loop with minimal delays (for human examples, see e.g. Brunnström et al., 2020; Caserman et al., 2019). There is, however, a lower limit on these delays when using conventional cameras and visual displays, as these have inherent latencies that are difficult to modify (Stowers et al., 2014). In addition, when recording tethered flight, at least one full wing stroke needs to be captured by each video frame, so wing movements need to be captured before stroke amplitude can be estimated, leading to a delay of at least one wing stroke period. When measuring walking behavior on a trackball, the ball’s motion can theoretically be recorded at high temporal frequency and instantly fed into the software controlling visual scenery. In practice, the trackball is often filmed at no more than 200 Hz (Bagheri et al., 2022; Dahmen et al., 2017).

One of our development aims was to combine several robust and easily accessed software packages to produce a flexible VR with short latencies. While MultiMoVR (Kaushik et al., 2020) was developed to reduce costs and uses the open source Panda3D game engine (Goslin and Mine, 2004), it uses Kinefly (Maimon et al., 2010) for tracking wing movements, which is sensitive to camera settings and light fluctuations. Recent advances in machine learning, and especially the development of DeepLabCut (DLC, Mathis et al., 2018), provides an alternative, efficient and robust method for tracking the WBA (Salem et al., 2022). Indeed, DLC has revolutionized behavioral neuroscience, as it allows for markerless pose estimation after training on relatively small data sets. FicTrac, which is often used for trackball experiments (e.g. Fenk et al., 2022; Loesche and Reiser, 2021; Turner et al., 2022), uses open-source software for reconstructing the fictive path of an animal walking on a patterned sphere filmed with a USB camera (Moore et al., 2014).

Unity (Unity Technologies), which has been used in VR arenas such as the Antarium (Kócsi et al., 2020), provides a user-friendly environment for creating virtual 3D scenes, and is more robust that Panda3D, with a larger community and better support. Unity is relatively easy to learn, and has been used for creating immersive learning environments (Needle et al., 2022) and for studying peripheral visual field loss in humans, as it provides proper perspective correction (Doyon and Jung, 2023).

We here show that the loop can be effectively closed using the Unity game engine together with machine vision (see also Müller et al., 2023). We used DLC-live (Kane et al., 2020) for tethered flight in hoverflies, and FicTrac (Moore et al., 2014) for trackball experiments in crabs, with delays below 50 ms. We validated the DLC-live-Unity connection by letting hoverflies navigate towards flowers, and the FicTrac-Unity integration by letting tethered crabs evading virtual crabs or birds. We also developed a Unity interface, CAVE, that performs all calculations based on markerless pose estimation input from DLC-live. CAVE allows the user to define the gain between the WBA and the tethered animal avatar’s yaw and thrust motion through the virtual world, and provides the ability to design experiments, and save data with little or no code.

## Materials and Methods

### Hardware and software

All hardware and commercial software for the tethered flight (Fig. 1A) and trackball arena (Fig. 2A) can be found in Table 2. In the tethered flight arena (Fig. 1A) each of the three screens subtended 118° x 142° (width x height) of the visual field. In the trackball configuration (Fig. 2A) each of the four screens subtended 90° x 49° of the visual field.

**Figure 1.**
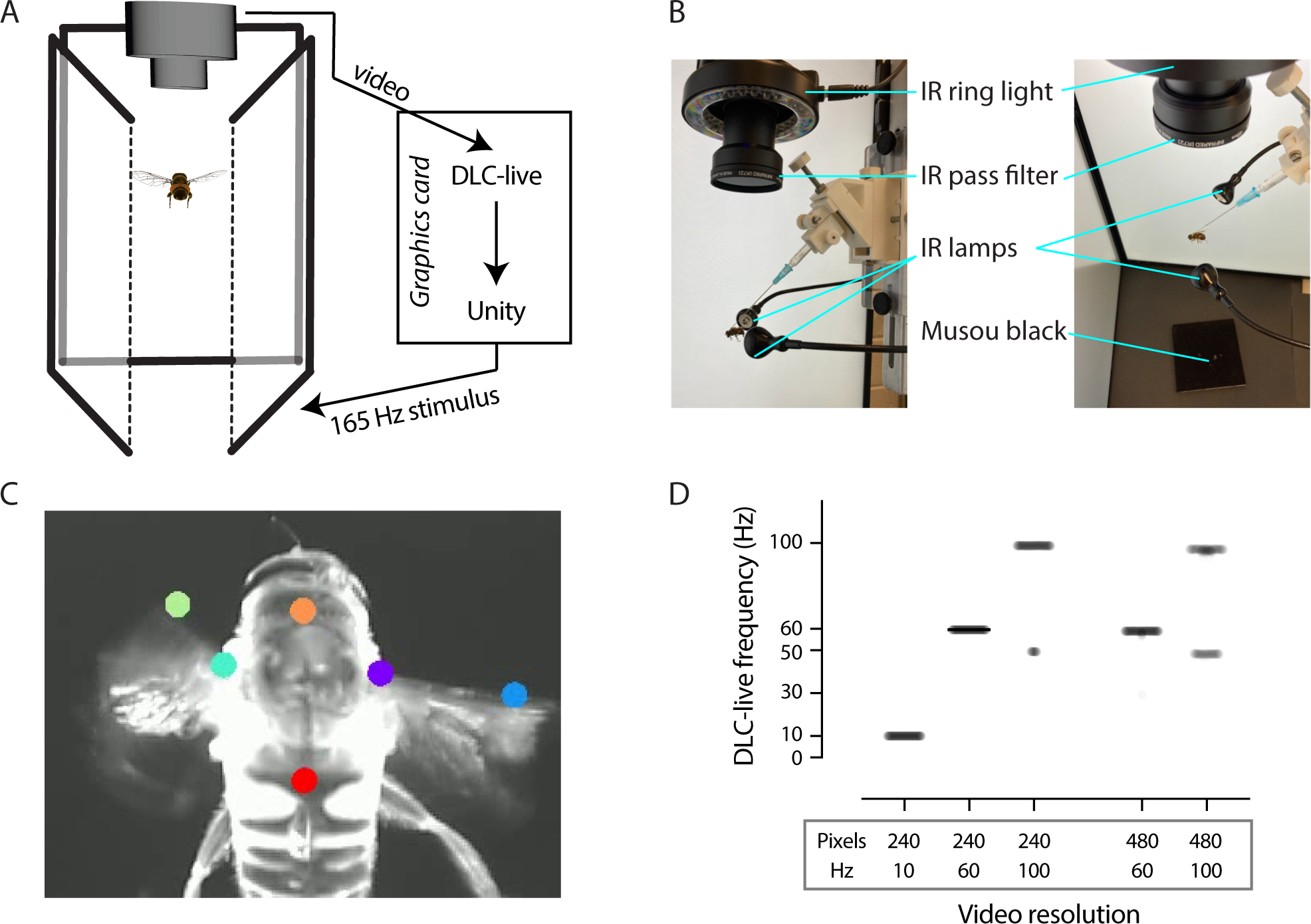
Virtual reality arena overview. A) The tethered insect (here a female *Eristalis tenax* hoverfly) placed in the center of three gaming monitors, oriented vertically. The insect is filmed from above using a PS3 camera (Kaushik et al., 2020) with the video fed into DeepLabCut-live (DLC-live) which performs markerless pose estimation in real time (Kane et al., 2020). Unity uses this information to calculate the requisite scene translation and rotation based on the wing beat amplitude and updates the virtual world displayed at 165 Hz. Hoverfly pictogram by Malin Thyselius. B) Lateral (left) and dorsal view (right) of the tethered *E. tenax* illuminated in infrared with two USB lights from the side, and an infrared ring light from above. The PS3 camera is equipped with an infrared pass filter, and a musou black surface is placed below the insect. C) Example video frame with two points on the anterior edge of each wing stroke and two points along the thorax tracked using DLC-live (Kane et al., 2020). D) The frame rate of the resulting DLC-live data using different spatial (240 x 320 or 480 x 640 pixels) and temporal (10, 60 or 100 Hz) video resolutions. Each semi-transparent data point shows the delay between two subsequent frame updates (N = 132, 427, 588, 483, 597).

**Figure 2.**
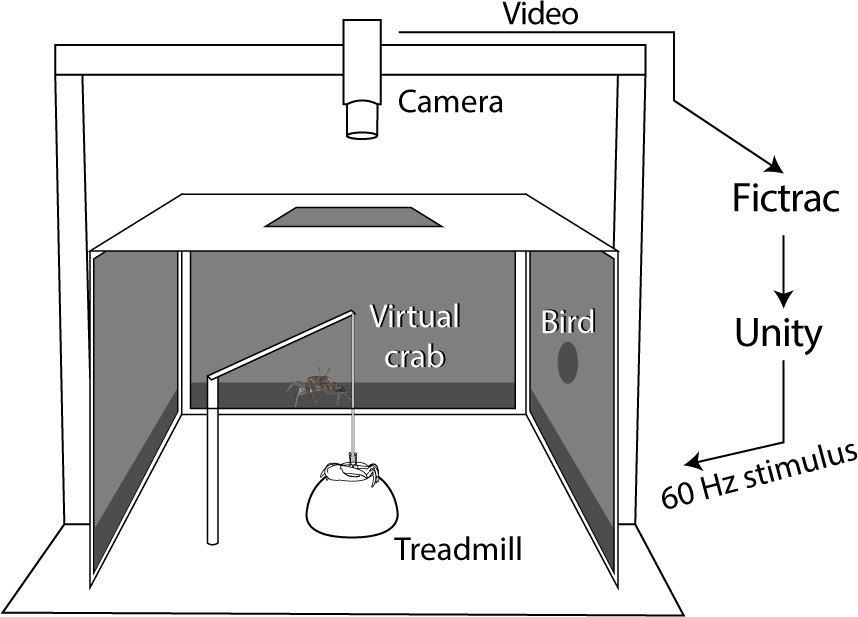
Trackball set-up. The crabs were placed on an air cushioned treadmill. Stimuli consisted of a virtual crab below the horizon (dark grey) and virtual bird above the horizon (pale grey) which were presented on either of the four adjacent stimulus monitors surrounding the treadmill. The experimental crab was held in place by a rod attached to its carapace that allowed it to slide up and down and rotate freely around the center of the ball, but not move away. The crab and the trackball were filmed from above using a FLIR Grasshopper3 camera with the video fed into FicTrac (Moore et al., 2014) which calculates ball’s motion in real time. Unity uses this information to calculate the requisite scene translation at the monitor update frequency (60 Hz).

**Table 1.**
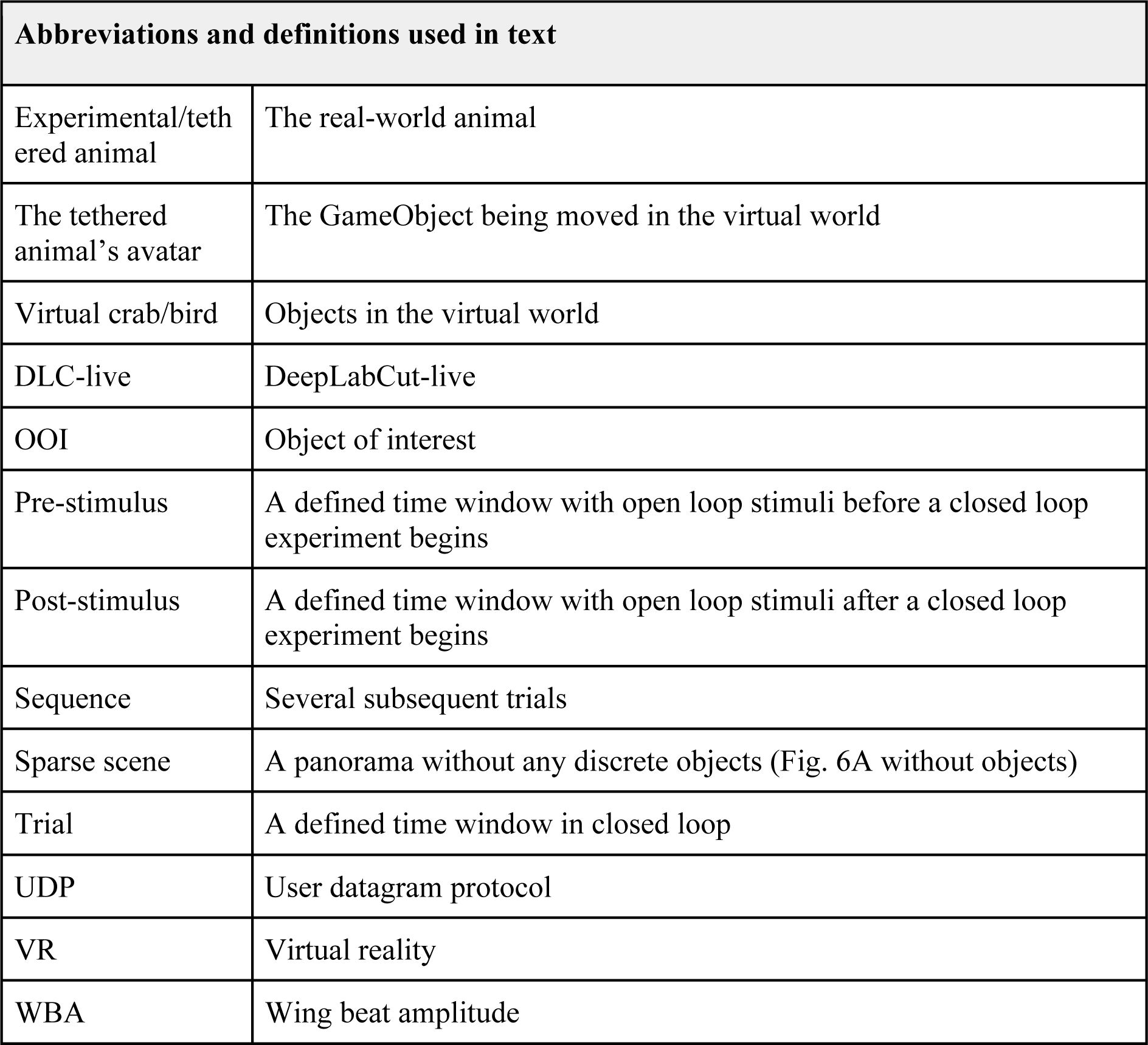
Abbreviations and definitions.

**Table 2.**
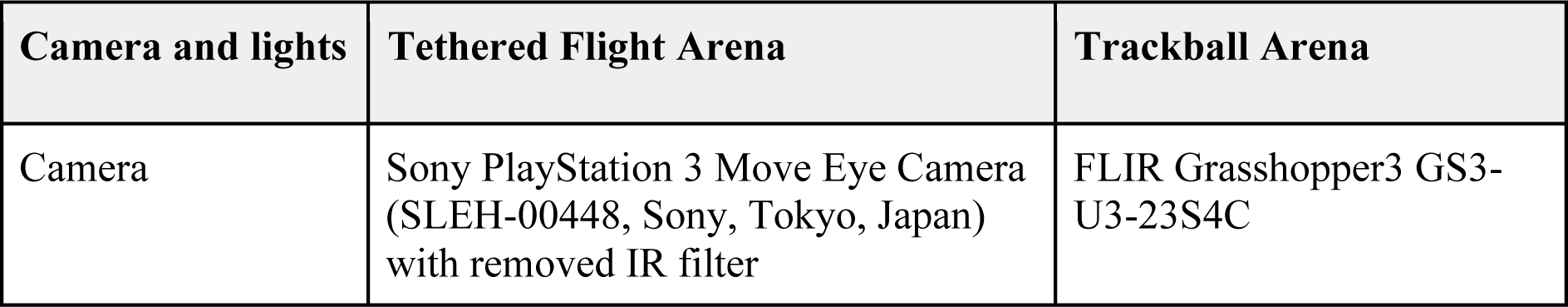
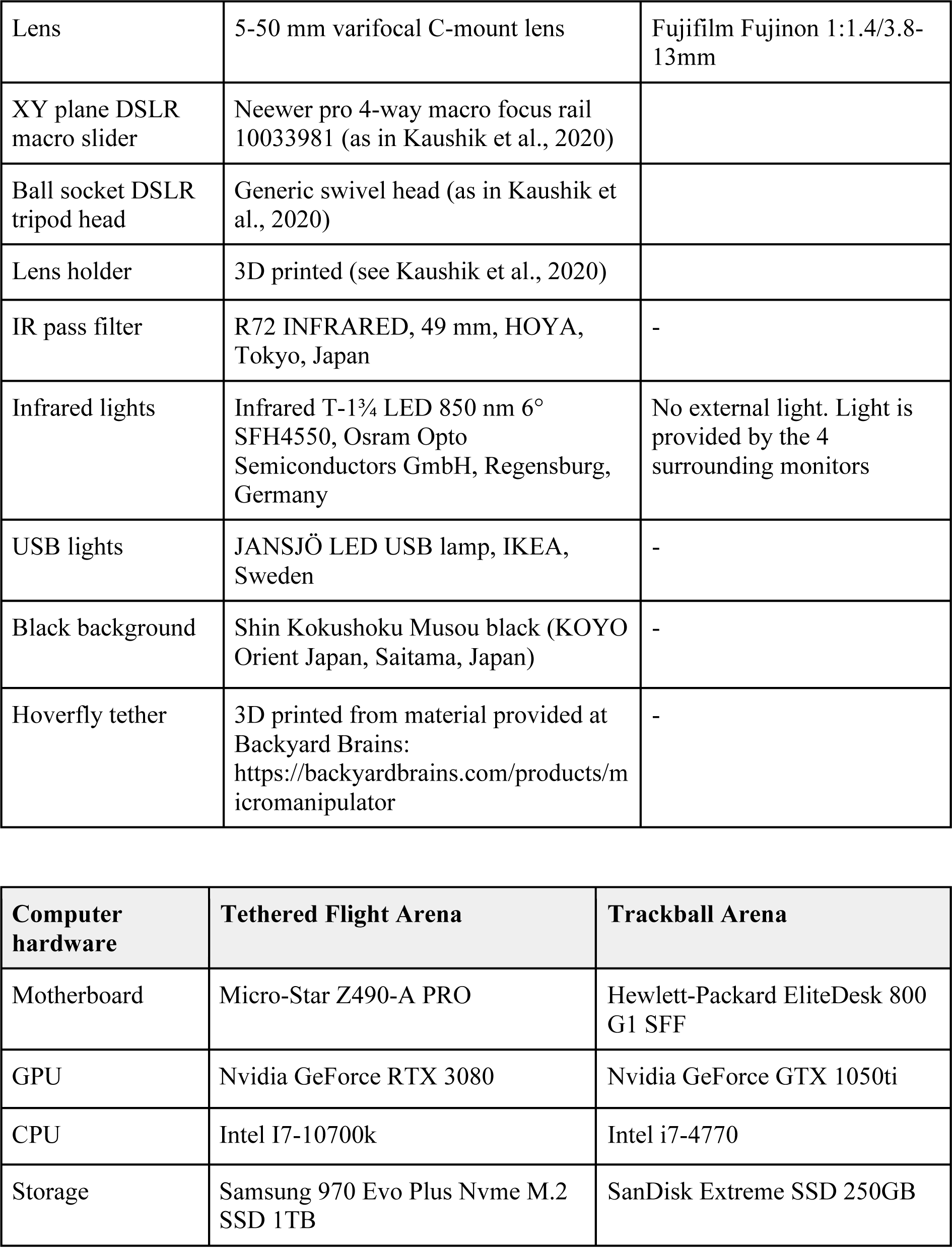
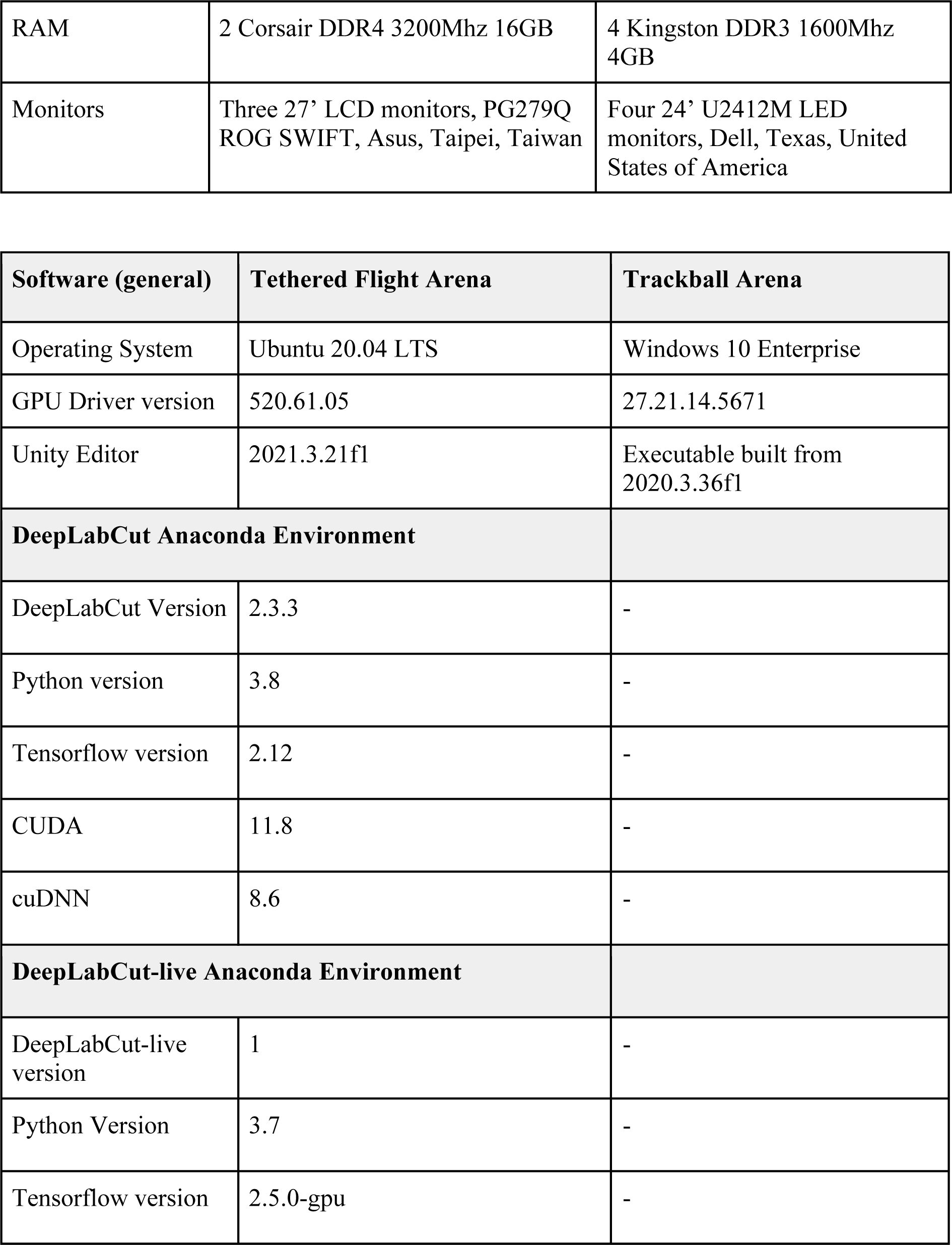
Hardware and software.

### Animals

For behavioral testing of the tethered flight configuration we used *Eristalis tenax* females, 14-24 days old, reared and housed as described previously (Nicholas et al., 2018). The hoverfly was tethered to a needle (BD Microlance 23G x 1 1/4” - 0.6 x 30 mm Blue hypodermic needles) at a 32° angle using a bee’s wax and resin mixture. The needle was then attached to a syringe (BD tuberculin syringe, 1 ml). To encourage flying we provided airflow manually. Once the hoverfly flew consistently, we placed it in the center of the arena (Fig. 1A, B) at 10 cm distance from the center of each screen. We kept encouraging flight using a sparse scene rotating at 0.33 Hz in open loop.

For behavioral testing of the trackball configuration we used the fiddler crab *Gelasimus dampieri* collected from intertidal mudflats near Broome (-17.9°S, 122.2°E), Western Australia. Crabs were housed in an artificial mudflat at the University of Western Australia and exposed to a tidal cycle of seawater inundation. They had a 12-hour light-dark cycle and their diet was supplemented with crab food. Crabs were treated according to UWA Animal Ethics Committee (AEC) approved methods (UWA AEC project number RA/3/100/1515). At experimental time, crabs were tethered to a sliding carbon-fiber rod by a magnet glued (Loctite Superglue, ethyl cyanoacrylate) to the crab’s carapace. They were then placed on an air-cushioned polystyrene treadmill ball (Bagheri et al., 2022; Donohue et al., 2022). The crabs were acclimatized (via video monitoring) until they began walking or feeding from the trackball before experimental stimuli were initiated.

### DLC model

We filmed tethered male and female *Eristalis tenax* responding to open loop stimuli in the tethered flight arena (Fig. 1A), including a sinusoidal grating (wavelength 35 mm, 5 Hz), a starfield stimulus (yaw at 50°/s), and a bar (width 5.5 mm, varying in height from 1.4 mm to 60 cm). We used DeepLabCut (DLC) version 2.3.3 (Table 2, and see Mathis et al., 2018) to train a model to track the thorax, and the peak downstroke angle, also referred to as the wing beat amplitude (WBA), of each wing (see e.g. Maimon et al., 2010). For this, we manually labelled the following six locations: tegula and tip of the left and right forewing, anterior thorax and anterior abdomen (Fig. 1C), for 16 extracted video frames each from videos of four individual animals (2 males, 2 females). These frames included examples of yaw rotation, forward translation and no flight. We trained the DLC model using 300,000 maximum iterations. The evaluated train error and test error were 1.2 pixels and 1.16 pixels, respectively.

The resulting model was used in DLC-live version 1 (Table 2, and see Kane et al., 2020), to track the same six points in real-time. We quantified DLC-live’s ability to perform markerless pose estimation of every unique frame under different video resolutions (Fig. 1D) by filming at two spatial resolutions (240 x 320 pixels and 480 x 640 pixels) and three temporal resolutions (10, 60 and 100 Hz, all below the wingbeat frequency of *Eristalis* hoverflies (Walker et al., 2010)). We quantified the time between sequential unique frames analyzed by DLC-live (“Time Frame was Captured” column in CAVE’s DLC-Data file, Fig. S1F), and converted this to temporal frequency.

### Unity and the CAVE interface

In the FicTrac-Unity configuration, the crab was attached to a magnetic tether, allowing it to rotate freely along the yaw axis (Fig. 2A, and see Donohue et al., 2022). The ball’s rotations in radians, obtained from the video recording of the ball (Supp Movie 1), multiplied by the ball’s radius, were used to update the crab’s avatar’s sideslip and forward motion (Moore et al., 2014) in the Unity generated virtual world. Jitter from FicTrac ball tracking noise was reduced by suppressing movements below a minimum distance over time, which was determined visually during preliminary testing. Rapid, random movement from tracking error was similarly limited by suppressing movements above a maximum distance. Because FicTrac errors are not cumulative, these adjustments did not affect overall tracking accuracy. Videos were filmed at a resolution of 544 x 638 pixels at 60 Hz.

In the DLC-live-Unity configuration we used the x-y-coordinates of the six points tracked in each frame (Fig. 1C) to calculate the WBA and the resulting yaw and thrust values for the avatar controller based on the gain information from the settings manager, available in our CAVE interface (Fig. 3, S1). This is described in the Results section.

**Figure 3.**
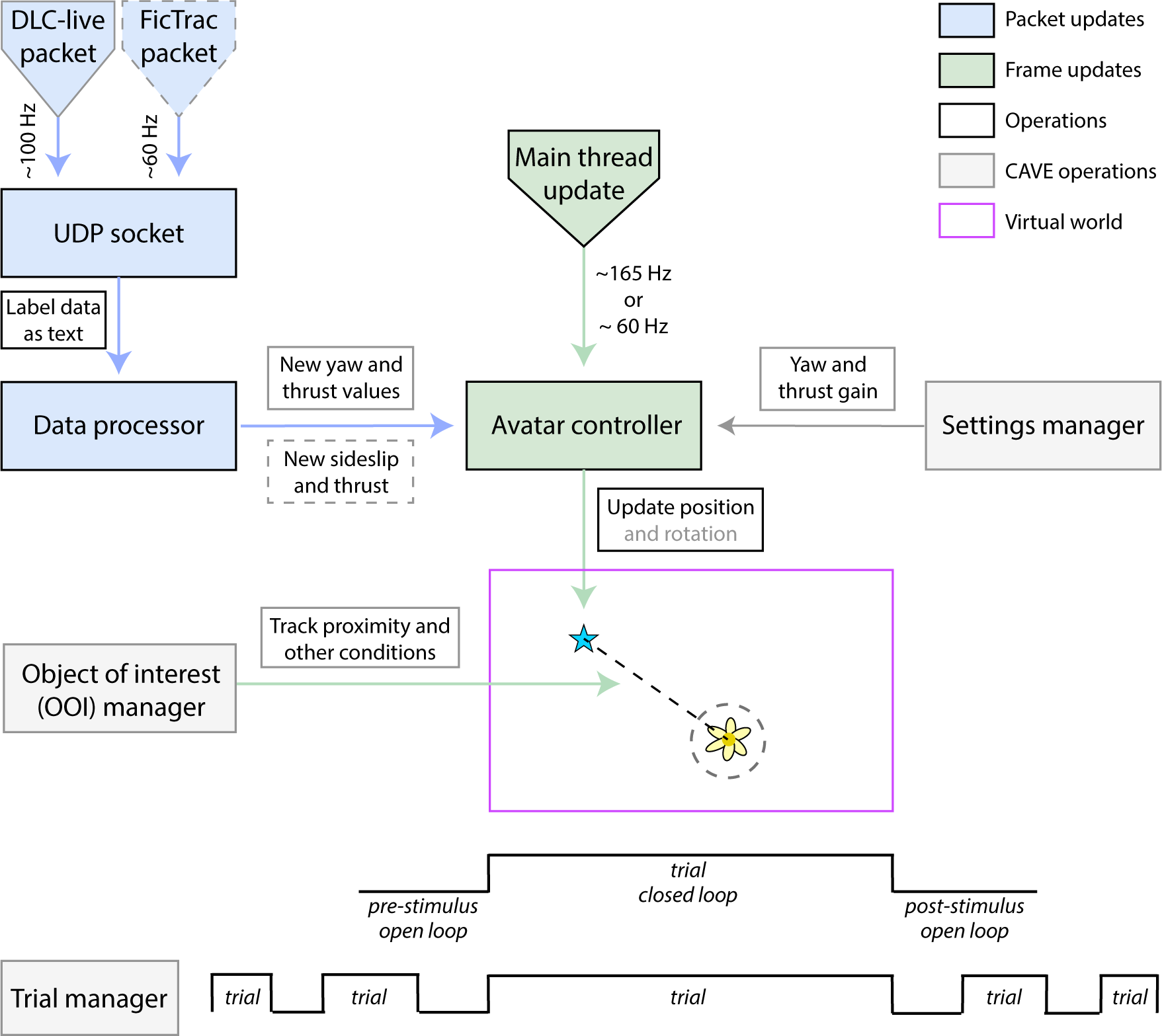
Technical diagram. The information flow from DLC-live (filmed at 100 Hz) or FicTrac (60 Hz) through the UDB socket and the data processor to the avatar controller. This updates the avatar’s new rotation (hoverfly only) and/or position (hoverfly and crab). In CAVE, this uses information from the settings manager. The main thread update runs at the stimulus display monitor frequency. The OOI manager tracks proximity (dashed circle) and other conditions between the avatar (blue star) and other OOIs (e.g., flower). Each trial (pink) is run in closed loop and is flanked by pre- and post-stimulus open loop times. The trial manager can be used to combine and interpolate trials.

In both cases, we used UDP sockets to receive the x-y-coordinates of the points tracked by DLC-live and their confidence, or the forward and sideward rotation of the treadmill ball provided by FicTrac (blue, Fig. 3, and see Aoukar, 2021). Upon receiving a packet, the C# script responsible for controlling the tethered animal’s avatar updates its rotation and/or position (green, Fig. 3), and the requisite main thread updates at the refresh rate of the screens (60 Hz for the crabs, 165 Hz for the hoverflies, Fig. 1A, 2A). In the tethered flight configuration, the stimulus screens were displayed at 165 Hz, and the video was filmed at 100 Hz. If there was no new DLC-live packet available, the avatar would keep performing thrust and yaw motion as defined by the previous packet (Fig. 3).

In CAVE, the object of interest (OOI) manager tracks the conditions of objects of interest, such as the distance to the tethered animal’s avatar (dashed line, Fig. 3). The trial manager is used to define trials, which contain the closed loop component, flanked by pre- and post-stimulus intervals in open loop (Fig. 3). The sequence manager is used to combine trials, and to define the open loop stimulus shown in between (Fig. 3, S1). The usability and testing of the CAVE interface is described in the Results.

### Behavioral validation, trackball

For behavioral testing of the trackball configuration, we exposed six fiddler crabs (*Gelasimus dampieri)* to two virtual objects: a crab walking on the ground and a flying bird. The crab was placed near a burrow 30 cm from the tethered crab’s avatar. The burrow was centered on one of the four screens. The crab moved in a cyclical fashion 10 cm away and then back to the burrow, spending 9 – 12 s inside the burrow and 21 – 35 s above it, depending on the movement speed. Movement direction and speed (0.75 - 1.25 cm/s) were randomized. In between cycles the virtual crab briefly descended into the burrow. The virtual bird, represented by a 3 cm diameter black sphere, appeared randomly on one of four surrounding screens, with the virtual crab randomly positioned either on the screen to the left or right of the bird. The bird could move in three different ways, but always started 200 cm from the avatar and 15° above the avatar’s visual horizon (Fig. 2B). During the “threatening” condition, 1) the virtual bird approached the crab in a straight line with a velocity of 19.9 cm/s from the start position and stopped moving at a virtual distance of 1.5 cm from the position the crab’s avatar was in at the beginning of the movement. In the two “non-threatening” conditions, the bird was either 2) stationary at the start position or 3) moved forwards and backwards along a 30° arc around the start position without coming closer.

We displayed these stimuli in either open or closed loop. In open loop, the crabs’ movements did not influence the appearance of the virtual stimuli on screen, whereas in closed loop, the visual scene adjusted based on the crabs’ position in the scene. Each trial was 60 s long. Both the virtual crab and bird were stationary at the start. After 5 s the bird started moving, and it kept moving for 12 s. To ensure adequate recovery time, there was a pause of at least 5 minutes between successive trials. We conducted six trials for each crab, presenting each bird movement condition with the virtual crabs positioned on both the left and right sides. The order of these trials was randomized using 3 x 3 Latin squares to prevent order bias.

### Behavioral validation, tethered flight

For behavioral validation of tethered flight, we used a sparse scene with two dandelion plants equidistantly placed 2 m from the hoverfly’s start position. Each plant was approximately 20 cm tall and had a 20 cm diameter with the leaves. We used a proximity criterion to complete each trial, defined as a 3D sphere of 40 cm radius around either dandelion for at least 1 s. The hoverfly’s avatar was programmed to fly 30 cm above the ground. At this height, the proximity radius in the horizontal plane is 26 cm. Each trial had a maximum duration of 30 s. As control, we used the same scene without any dandelion plants, and a 30 s duration. In six females we performed control (C) and dandelion (D) trials in the following order: C-C-C-C-C-C-D-D-D-D-D-C-C-D-D-D-D-D-C-C-D-D-D-D-D-C-C-D-D-D-D-D-C. The first six control trials were used for data analysis, whereas the interspersed control trials were used to control for learning and fatigue. In between each trial we showed the sparse scene rotating at 0.33 Hz in open loop. In addition, we excluded trials where the hoverfly stopped flying.

The proximity rate was defined as the ratio of trials where the tethered hoverfly’s avatar triggered the proximity criterion. The total path length (*L*) for each trial was quantified as the vector sum of the tethered hoverfly avatar’s location changes. Path straightness was defined as *D/L* (Benhamou, 2004), where *D* was the distance between the start and end position of the hoverfly’s avatar in each trial, and *L* was the total path length for trials ended fulfilling the proximity criteria.

### CAVE data extraction and latency measurements

We extracted the WBA difference (WBAD, defined as |WBA_Left_ – WBA_Right_|) and the WBA sum (WBAS, defined as WBA_Left_ + WBA_Right_) from the DLC-data.csv file (Fig. S1F). We extracted the resulting avatar yaw and position using the y-rotation and x-z coordinates reported in the Transform.csv file (Fig. S1F).

We used photodiodes to determine the time for closing the loop, using a video with a square flickering between black and white, played on an iPhone placed in the hoverfly’s position (Fig. 1A). This resulted in a corresponding black-to-white change on the stimulus screens. The black-to-white change on both the iPhone and the stimulus screen were recorded with photodiodes at 10 kHz. This latency was compared to the data reported in CAVE’s DLC-Data.csv file (Fig. S1F). The time between each video frame update was defined as the difference between sequential unique instances of “Time Frame was Captured”. The time taken by DLC-live to perform its pose estimation is reported in the “DLC_Latency” column in the DLC-Data.csv file (Fig. S1F). To quantify the time taken by Unity to calculate the WBA and the requisite scene update, we used the following equations based on information from the DLC-Data.csv file (Fig. S1F):

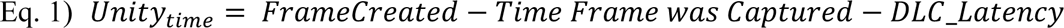

We quantified the time to close the FicTrac-Unity loop using a mirror and camera, allowing the ball’s movements and the subsequent screen updates of a vertical line to be recorded simultaneously. We used the cross-correlation between the pixel position of the vertical line on the screen reflected by the mirror and the rotation of the ball in the x-axis from FicTrac to estimate latency. The video was recorded at 60 Hz, giving a 16.7 ms resolution.

### Statistical analysis

Crab movement trajectories relative to the virtual crab and bird were analyzed in R (Core, 2008). Path coordinates were transformed such that virtual crabs were aligned at 0° and virtual birds at 90°. We then used linear mixed effects models using the lme4 package (Bates et al., 2015) to calculate the statistical significance of feedback condition (open or close loop) and bird actions on distances crabs travelled away from the virtual crab and bird over the duration of the experiment. Animal ID was included as a random effect. Significance (p < 0.05) of fixed factors was determined by comparing nested models that differed by only one factor. All p-values presented were estimated by comparison with the final model (using Likelihood Ratio Tests) that only contained significant fixed factors. The assumptions of the models were checked by exploring the distribution of the residuals (using Q–Q plots) and examining plots of the standardized residuals against the fitted values for each model.

Analysis of tethered flight data was performed in Matlab (R2021b, Mathworks) and Prism 10.1.0 for Mac OS X (GraphPad Software), with statistical test, p-values, and sample size indicated in the figure legend, where N refers to the number of animals, and n to the number of trials. As path straightness depends on the flight duration, we needed to correct for this in our statistical comparisons. As four control trials triggered the proximity criterion, we randomly selected four out of the other 32 control trials and cropped these to the same duration before statistical analysis. Similarly, as 36 dandelion trials triggered the proximity criterion, we randomly selected 36 out of the remaining 77 dandelion trials and cropped these to the same durations.

### Data and code availability

All data and analysis scripts have been submitted to DataDryad: Private DOI for peer review: https://datadryad.org/stash/share/vzNau2jRPU3JVEQlEqkLwNSu1bbZ3TBdG-d0X6881lA

The FicTrac-Unity interface can be found at Github: https://anonymous.4open.science/r/fiddlercrabvr-7CB1/README.md

The CAVE interface, user manuals, user videos and other documentation is via Github: https://anonymous.4open.science/r/2489/README.md

## Results

### A virtual reality arena using machine vision and Unity

We developed closed loop, virtual reality (VR), by combining machine vision with the Unity gaming engine (https://unity.com). For flying insects, we used MultiMoVR (Kaushik et al., 2020) as a foundation, but updated it using Unity and DLC-live (Kane et al., 2020; Mathis et al., 2018). We filmed the flying insect from above (here an *Eristalis tenax* hoverfly, Fig. 1A-C) using a PS3 camera (as in Kaushik et al., 2020) equipped with an infrared pass filter. The hoverfly was illuminated with infrared lights, with a musou black surface below maximizing the contrast (Fig. 1B).

We trained a DLC model (Kane et al., 2020; Nath et al., 2019) to track six points on the hoverfly (colored dots, Fig. 1C). Two of these were along its longitudinal axis, and two each along the anterior edge of each wing stroke (Fig. 1C). To determine the performance of DLC-live, we measured its update frequency under different video resolutions. We first fixed the spatial resolution at 240 x 320 pixels and found that at 10 and 60 Hz DLC-live runs reliably without missing any frames, but at 100 Hz 15% of the video frames were updated at 50 Hz (Fig. 1D). We next fixed the video’s spatial resolution at 480 x 640 pixels and found that at 60 Hz DLC-live runs reliably without missing any frames, and at 100 Hz 36% of the frames are run at 50 Hz (Fig. 1E). If high temporal fidelity is crucial, increasing the graphics card performance, the CPU, or reducing the temporal or spatial resolution of the video camera, can thus be advantageous (hardware specifications, Table 2). From here on we show tethered flight data after filming at 240 x 320 pixels at 100 Hz (Fig. 1D), which is below the wingbeat frequency (149 – 180 Hz) of *Eristalis* hoverflies (Walker et al., 2010).

For the trackball set-up, we recorded movement live at a resolution of 544 x 638 pixels at 60 Hz using a dorsally mounted camera and FicTrac (Fig. 2). During testing we chose this resolution and temporal frequency to allow both Unity and Fictrac to run on a single computer without skipping frames. Resolution and framerate were limited by the use of low-specification computer hardware (Table 2, Trackball arena column) and could easily be improved with a higher quality display, graphics card and CPU.

### Escape responses in crabs

To test the FicTrac-Unity closed loop (Fig. 2, 3), we used tethered fiddler crabs walking on a trackball and compared their responses to a virtual crab and bird in closed and open loop. In both open and closed loop scenarios, we found that the tethered crabs’ avatars ran significantly further away from the bird when it was looming (Fig. 4A, E; red, Fig. 4D; χ^2^ = 6.52; Df = 2; *P* = 0.0385, linear mixed model). The fast-moving nature of the looming virtual bird reduced the importance of the feedback and there was no difference in how far the crabs ran away from the looming bird between open and closed loop (red, Fig. 4D; χ^2^ = 1.25; Df = 1; *P* = 0.264, linear mixed model).

**Figure 4.**
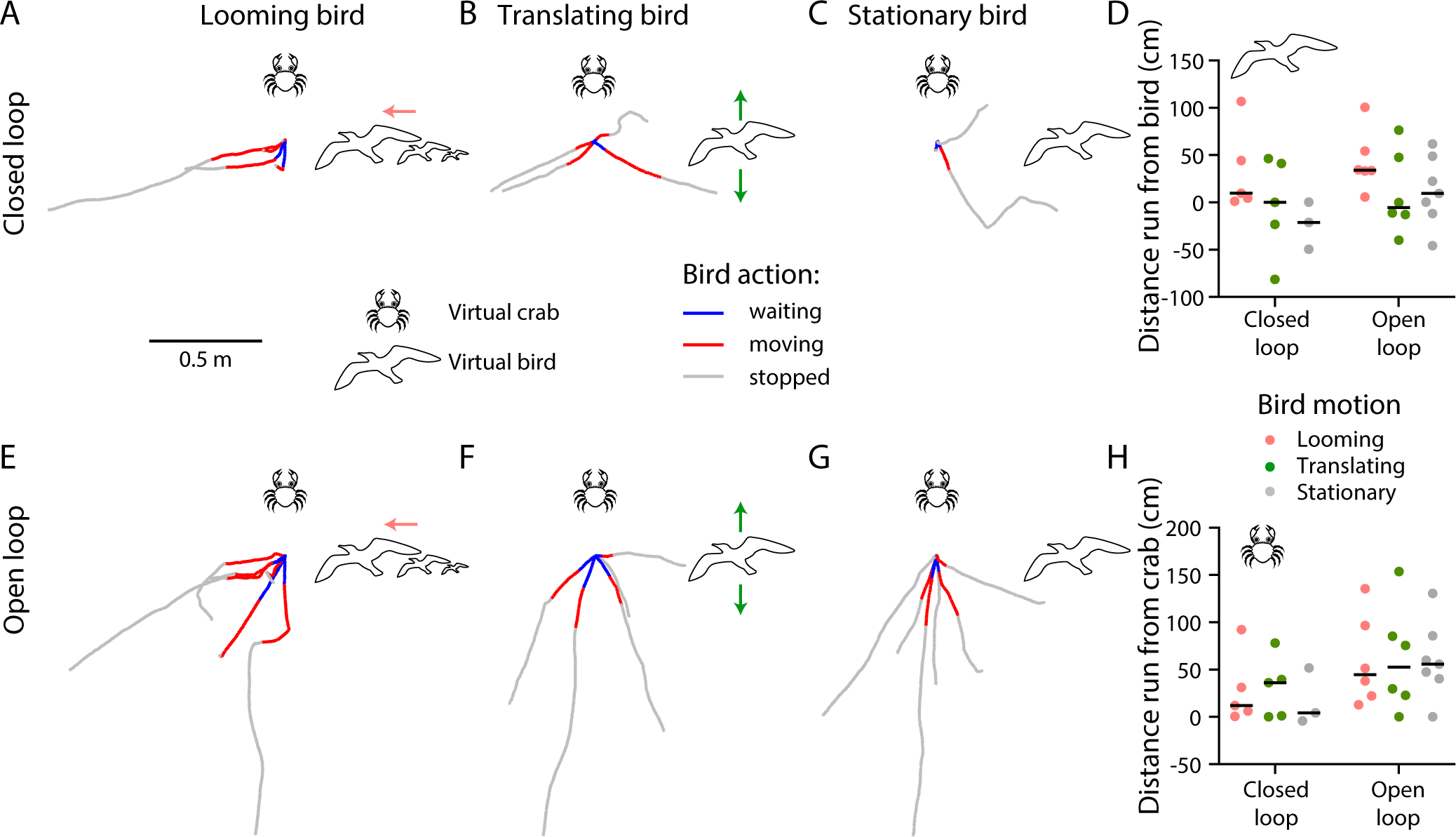
Tethered crabs walk further away from virtual crabs in open loop. A) The walking trajectory of a tethered fiddler crab on a trackball in closed loop, being exposed to a virtual crab 30 cm away and a virtual, looming bird at a starting distance of 200 cm (193 cm on the horizontal plane, not to scale). B) The walking trajectory of a tethered fiddler crab being exposed to a virtual crab and a translating bird. C) The walking trajectory of a tethered crab being exposed to a crab and bird. D) The distance run from the virtual bird in open (left) and closed loop (right), color coded according to the bird’s motion. E) The tethered fiddler crab walking trajectory in open loop, in response to a virtual, crab and a virtual, looming bird. F) The walking trajectory in response to a crab and a translating bird. G) The walking trajectory in response to a crab and bird. H) The distance run from the virtual crab in closed (left) and open loop (right), color coded according to the bird’s motion. In panels A-C and E-F, the data are color coded according to the bird’s actions.

However, the response to the virtual crab differed between open and closed loop. In closed loop (Fig. 4A-C), the experimental crab’s effort to move away from the virtual crab resulted in the virtual crab visually receding into the distance, accurately replicating the dynamics of a stationary object in the environment. In open loop, the experimental crab’s ran approximately twice as far from the virtual crab (Fig. 4H, χ^2^ = 4.29; Df = 1; *P* = 0.0383, linear mixed model), irrespective of the virtual bird’s behavior (Fig. 4; χ^2^ = 0.213; Df = 2; *P* = 0.899, linear mixed model). In open loop the tethered crabs could not increase their distance to the virtual crab, which, consequently, would have appeared to follow the experimental crab as it moved (Fig. 4E-H). The experimental crabs moved away from the virtual crab even when there was a larger threat of a looming bird (Fig. 4E). Our data (Fig. 4) thus suggest that closed loop experiments offer a more accurate simulation of non-threatening stimuli in environments.

### CAVE user interface developed for tethered flight

For tethered flight VR, the six points (Fig. 1C) were used to calculate the left and right wingbeat amplitude (WBA_L_ and WBA_R_, Fig. 5A, E), relative to the longitudinal axis (black line, Fig. 5A, E). Note that it is important that DLC-live tracks the two points underlying the longitudinal axis robustly, as this is used as a ground truth (black line, Fig. 5A, E). We assume that if one wing has a larger WBA than the other (Fig. 5A), this represents an attempted yaw turn in the opposite direction (Maimon et al., 2010). We developed a Unity user interface for tethered flight, CAVE, that allows user control (grey, Fig. 3). In the CAVE settings manager (grey, Fig. 3), the user can define the relationship between the WBA difference (WBAD, defined as |WBA_L_ – WBA_R_|) and the resulting yaw (Fig. 5B), which has an inherent trade-off between maneuverability and stability (Kaushik et al., 2020). E.g., if the gain is high, the insect can turn rapidly, but this may result in looping flight patterns. If the user wants WBAD above or below a certain value to have less effect on the resulting yaw, the variable setting can be used (Fig. 5C, D).

**Figure 5.**
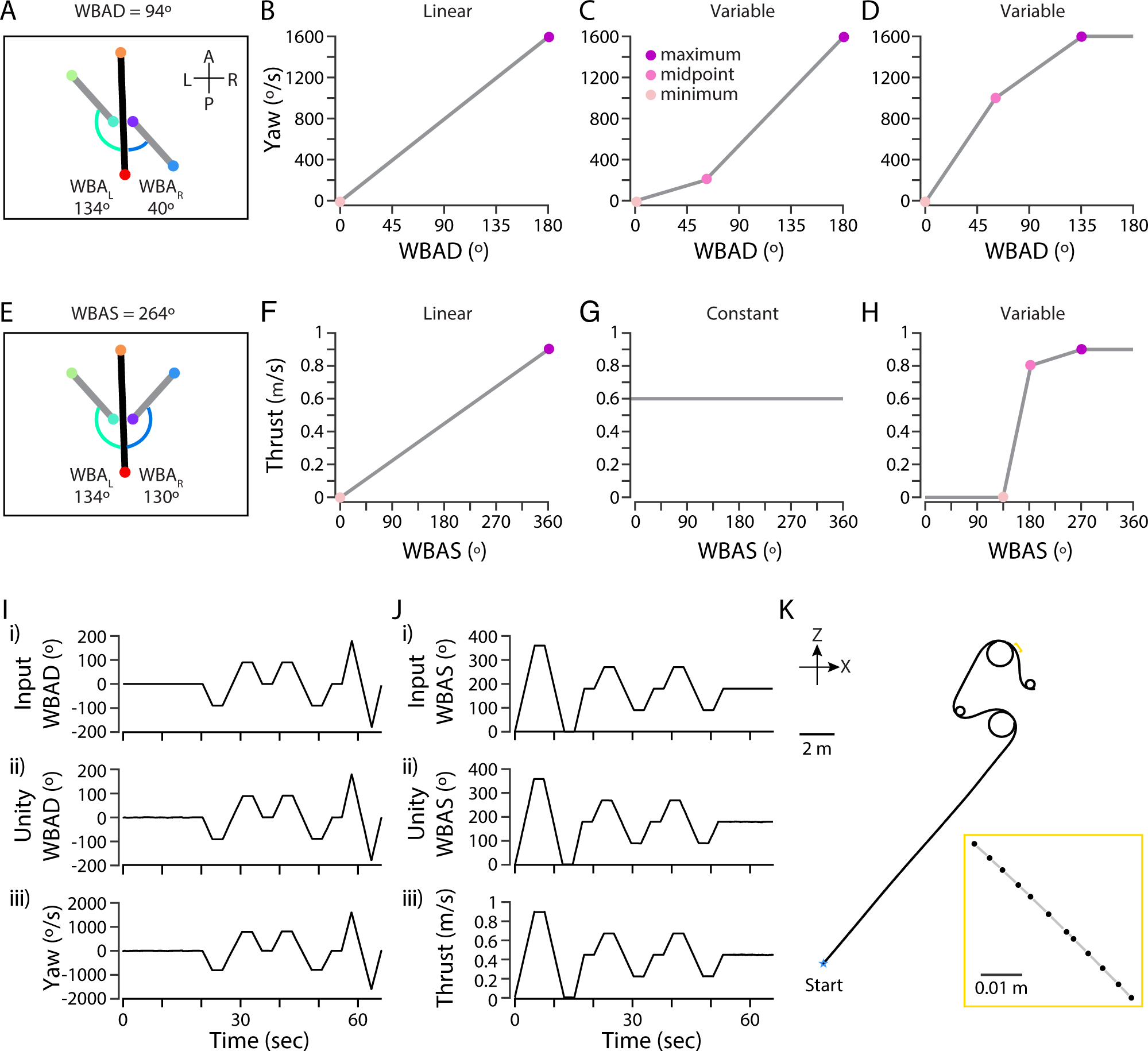
Yaw and thrust calculation in Unity. A) The wingbeat amplitude of the left (WBA_L_) and the right (WBA_R_) wing, with the tracked points color coded as in figure 1C. The WBA difference (WBAD) was defined as |WBA_L_ - WBA_R_|. B) With the linear setting the WBAD is directly proportional to the yaw. C) Using the variable setting the user defines the maximum WBAD and yaw, the midpoint WBAD and yaw, and the minimum WBAD and yaw. D) If the WBAD is above the defined maximum, the function plateaus. E) The WBA sum (WBAS), defined as WBA_L_ + WBA_R_. F) With the linear setting the WBAS is directly proportional to forward thrust. G) The forward thrust set to constant. H) If the WBAS is below the defined minimum, the function plateaus. I) *i)* The WBAD in our control video (Supp Movie 2) portraying a model (as in panels A, E) as a function of time. *ii)* The WBAD calculated by Unity, using DLC-live information. *iii)* The resulting yaw rotation performed by the avatar in the simulated environment using the linear setting in panel B. J) *i)* The WBAS in our control video (Supp Movie 2) as a function of time. *ii)* The WBAS calculated by Unity. *iii)* The resulting thrust by the avatar using the linear setting in panel F. K) The resulting trajectory (from panels I*iii*, J*iii*) as seen from above.

We used the sum of the left and the right WBA (WBAS, Fig. 5E) to define the tethered animal avatar’s forward thrust (Fig. 3) as a linear (Fig. 5F) or variable (Fig. 5H) relationship. Alternatively, thrust can be constant (Fig. 5G). The gains between WBA and avatar motion should ideally replicate the flight dynamics of the species investigated, to allow for naturalistic behavior (Kaushik et al., 2020).

To validate that CAVE executes the yaw and thrust calculations for the avatar movement in the virtual world correctly we created a video (Supp Movie 2) where we varied the WBA_L_ and the WBA_R_ of a stylized insect (as in Fig. 5A, E). The graphs show the resulting WBAD (Fig. 5I *i*) and WBAS (Fig. 5J *i*) as a function of time. The WBAD and the WBAS as extracted by CAVE (Fig. 5I, J *ii*), and the resulting yaw (Fig. 5I *iii*) and forward thrust (Fig. 5J *iii*) of the avatar, after using the linear settings described above (Fig. 5B, F), confirm that this was done correctly. The resulting trajectory seen from above (Fig. 5K) highlights that due to the mismatch in sampling frequency between the input video (100 Hz), and the display monitors (165 Hz), the avatar’s positions are not always evenly spaced (inset, Fig. 5K).

### CAVE experimental design

CAVE allows the user to set-up experiments (grey, Fig. 3, S1). Before each experiment, the user chooses a scene and populates it with objects. These can be scaled as desired (here scaled to have the same size, Fig. 6A, B), and defined as objects of interest (OOI, Fig. S1A), such as a stationary flower (yellow and red, Fig. 6A, B) or a moving insect or predator (brown and black, Fig. 6A, B). The user defines the initial OOI behavior, the encounter behavior and defines an encounter (Fig. S1A). For example, a flower can appear when the tethered animal’s avatar is within a certain distance, or a predator can follow the tethered animal’s avatar’s motion at a defined speed.

**Figure 6.**
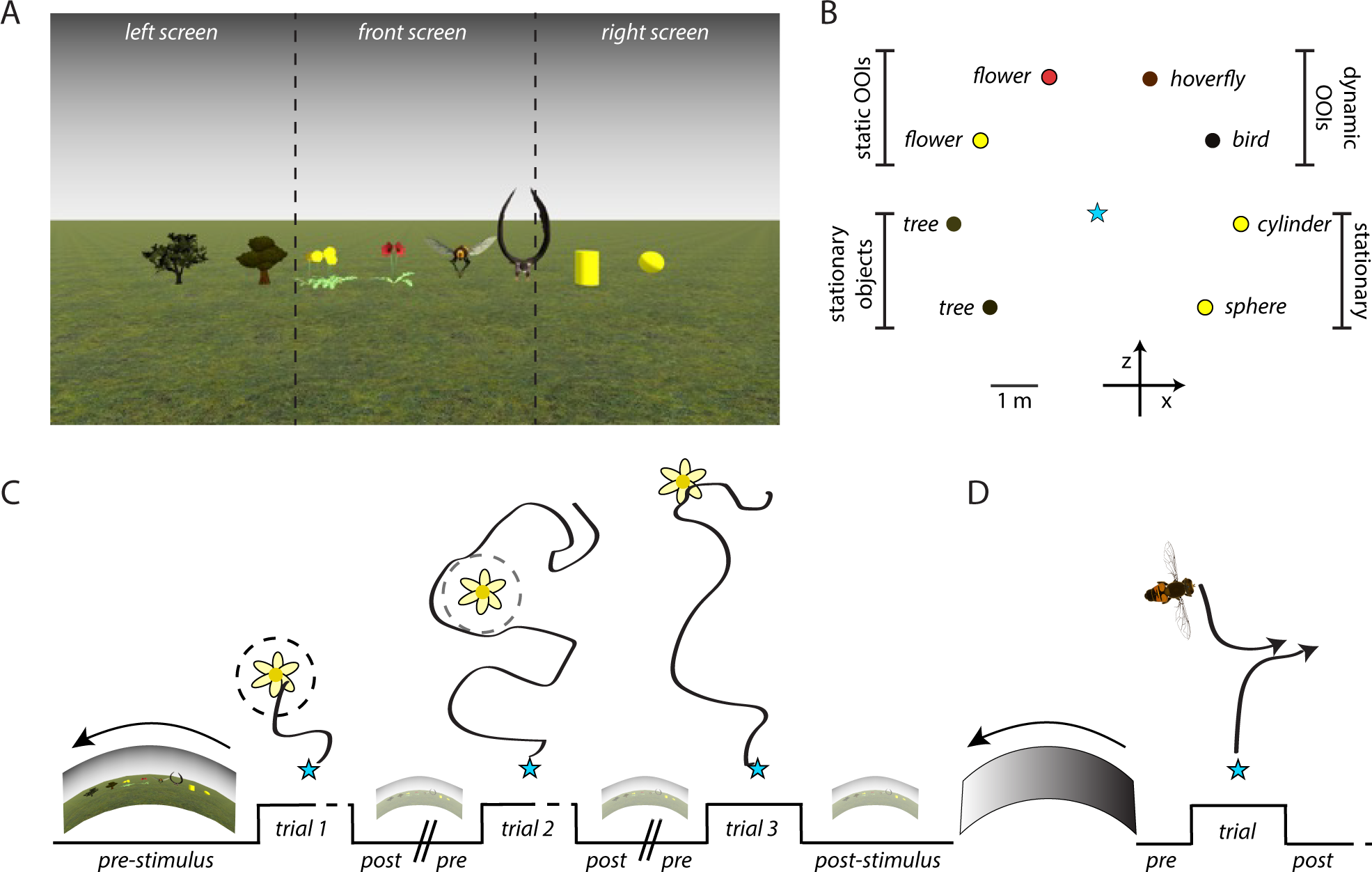
CAVE usability. A) An example sparse scene with eight objects placed equidistantly from the tethered insect’s avatar, scaled to have similar size. B) The same scene with the objects color coded as in panel A, seen from above. In this example the trees, cylinder and sphere are defined as objects, whereas the flowers, hoverfly and bird are defined as objects of interest (OOI). The turquoise star indicates the tethered insect’s avatar’s start position. C) An example sequence of 3 trials where the tethered insect’s avatar’s start position is indicated with a star, and a static OOI (a yellow flower) has been linearly interpolated to be placed at 3 different distances. Trial 1 and 2 use the proximity setting to end when either the proximity criterion (dashed circle around the flower, trial 1), or a duration (trial 2), have been fulfilled. Trial 3 ends after a fixed duration. During pre- and post-stimulus the scene rotates in open loop. D) An example sequence consisting of a single trial, using a dynamic OOI (a hoverfly) following the tethered insect’s avatar. In between the sequences a scene rotates in open loop.

Next, the user creates a trial, which is flanked by defined open loop pre- and post-stimulus times (Fig. 3, 6C, S1B), which can be used to habituate the fly to the virtual surround or entice flying using the optomotor response. The trials are combined with interventions to, for example, move an OOI to a new location if the tethered animal’s avatar is within a certain distance (Fig. S1B). Each trial ends when a proximity criterion is fulfilled or after a fixed duration. For example, here we have defined a proximity criterion as a 40 cm distance from the OOI (a dandelion, dashed circles, Fig. 3, 6C, S1B) for at least 1 s. In trial 1, the avatar fulfilled this criterion, whereas in trial 2 it was never close enough to the flower, and thus the trial continued for the defined duration (Fig. 6C). Trial 3 was set up to run for a fixed duration (Fig. 6C).

Trials can be put together into a sequence (Fig. 3, 6C, S1C). By using interpolations, an OOI can, for example, be placed at three evenly spaced distances within the trials of a sequence (flower, Fig. 6C). Several sequences can be put together before the experiment (Fig. 6D). The user defines what happens between sequences, in this example an open loop rotation (Fig. 6D). This open loop stimulus ends when the user presses *Next* to initiate the next sequence.

During the experiment (Fig. S1E), each trial’s data are continuously saved as lists. When a trial ends, the most recent lists are saved as CSV files (Fig. S1F), and the previous lists cleared to save memory. The Sequence and Trial files contain all experimental settings described above. The DLC-Data file contains frame-by-frame data from DLC-live (Fig. 3, S1F). The Transform files contain rotation and position data for the tethered hoverfly’s avatar (e.g. Fig. 5K), as well as for each OOI. This will, for example, include position data for dynamic OOIs (Fig. 6D).

### Hoverfly navigation

To test whether hoverflies navigate in the CAVE controlled VR, we used a sparse scene as well as one with two OOIs, each a yellow dandelion plant (Fig. 7A, B). We used linear settings for yaw and thrust (see Fig. 5B, F). We found that the hoverflies’ avatars flew towards the yellow dandelions, with a median proximity rate of 0.25 (dark pink, Fig. 7B, C, Supp Movie 3). When not fulfilling the proximity criterion, the trials lasted for 30 s (e.g. pale pink, Fig. 7B). In the control condition where no dandelions were visible, we analyzed post experiment whether the hoverflies would have fulfilled the proximity criterion (solid circle, dark green, Fig. 7A), and found a median proximity rate of 0.1667 (Fig. 7C, *P* = 0.0312, Wilcoxon matched-pairs sign ranked test).

**Figure 7.**
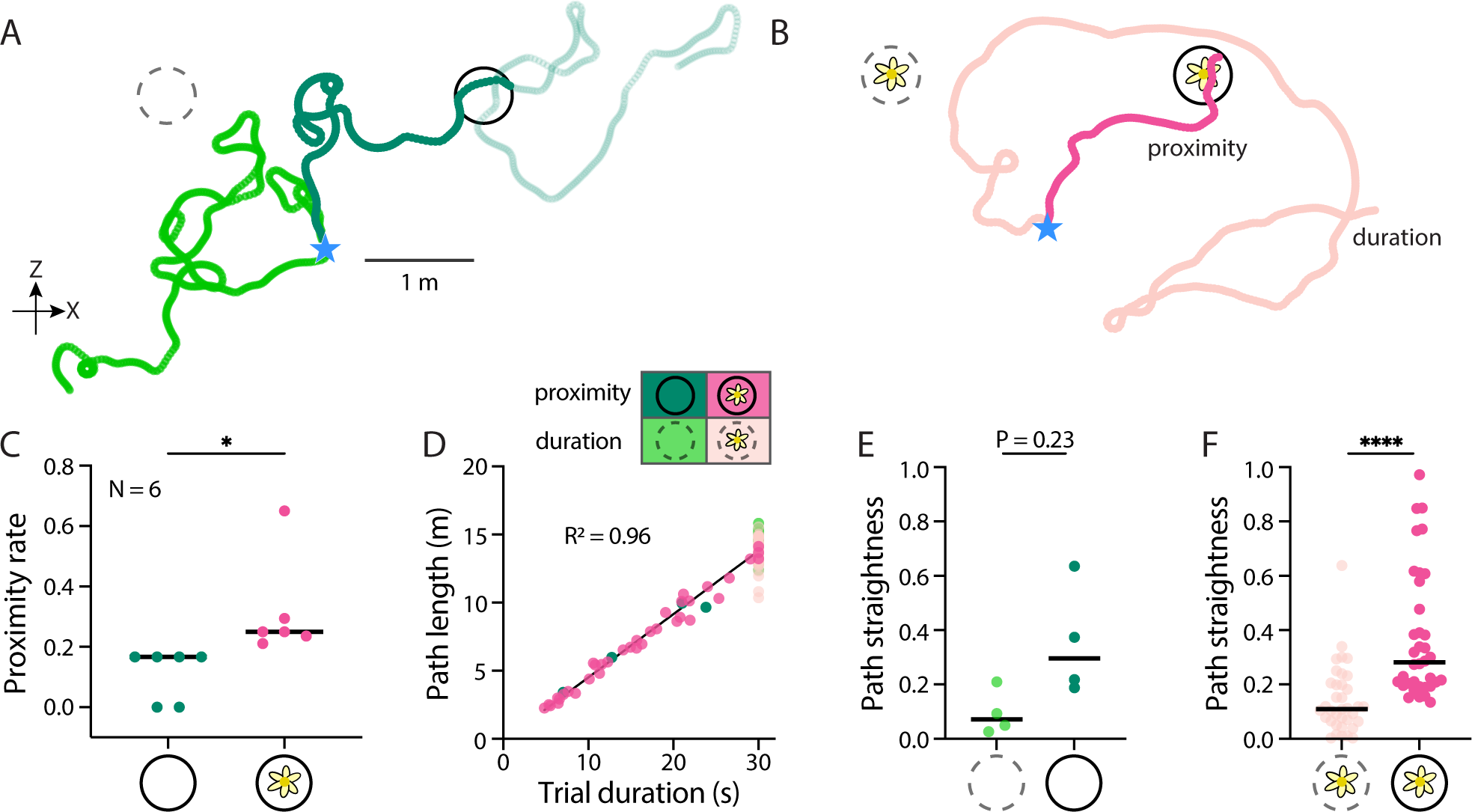
Hoverflies navigate within the virtual world. A) Two example flight paths of a tethered female *E. tenax* hoverfly’s avatar as seen from above (blue star indicates start position), with no OOI in the scene. We used the linear setting for yaw (see Fig. 5B) and thrust (see Fig. 5F), and the duration setting (30 s). Circles indicate the proximity size (26 cm horizontal radius at 30 cm height) used for proximity rate calculation post experiments. The dark green example would have triggered the proximity criterion (solid circle). The pale green data shows the flight path after this trigger. In contrast, the bright green example would not have triggered the proximity criterion. B) Two example flight paths of the same tethered female, with two OOIs (yellow dandelions) placed equidistantly from the start position (blue star). We used the same linear settings for yaw and thrust, and the proximity setting for ending each trial (26 cm horizontal radius, for 1 s, circles), with a total maximum duration of 30 s. The dark pink example triggered the proximity criterion (solid circle), while the pale pink did not (dashed circle). C) The proximity rate using the scene with no flowers or with two flowers (N = 6, Wilcoxon matched-pairs sign ranked test). D) Path lengths in the scene with no flowers (green), or in the scene with two flowers (pink), as a function of trial duration (simple linear regression test, R^2^ = 0.954). These have been separated into trials that triggered the proximity criterion (dark green and pink) and those that ended after 30 s duration (pale green and pink). E) Path straightness in the control scene, with the data color coded as in panel D (N = 6, n = 4, Kolmogorov-Smirnov test). F) Path straightness using the dandelion scene with the data color coded as in panel D (N = 6, n = 36, Kolmogorov-Smirnov test).

We quantified the path lengths from each trial as a function of trial duration. We found that the path length was directly proportional to the trial duration (Fig. 7D, R^2^ = 0.955, Simple linear regression test), indicating that the hoverflies did not modulate their flight speed a lot.

We expected the paths to be straighter if the flight towards the flowers was directed. To investigate this, we compared the path straightness when the proximity criterion was fulfilled (saturated colors, Fig. 7E, F) with trials where it was not (pale colors, Fig. 7E, F). We found that in the control scene, with no dandelions present, there was no significant difference between trials where the proximity criterion was triggered and those where it was not (Fig. 7E, *P* = 0.23, Kolmogorov-Smirnov test, n = 4). In contrast, in the dandelion trials the paths were significantly straighter when the proximity criterion was triggered compared with when it was not (median 0.1096 and 0.2810, respectively, Fig. 7F*, P* < 0.0001, Kolmogorov-Smirnov test, n = 36). Taken together, this suggests that the hoverflies were performing a directed flight toward the virtual dandelions when triggering the proximity setting.

### Closing the loop

We next investigated how long each step in the tethered flight loop-closing process takes (Fig. 1A, 8A). By using the DLC-live data reported in the DLC-Data file (Fig. S1F), we extracted the time between each new frame update from the video camera. As the camera had a refresh rate of 100 Hz, we expected this to be 10 ms. However, we found that 14% of the frames had a 20 ms delay, and 0.8% a 30 ms delay (11.5 ± 3.8 ms, mean ± std, “Video”, Fig. 8B, see also Fig. 1D). It took DLC-live 15.9 ± 3.4 ms (mean ± std, Fig. 8B) to perform the markerless pose estimation of the six points (Fig. 1C). Unity used 3.7 ± 2.0 ms (mean ± std, Fig. 8B) to calculate the WBA, and the resulting yaw and thrust movements of the avatar.

**Figure 8.**
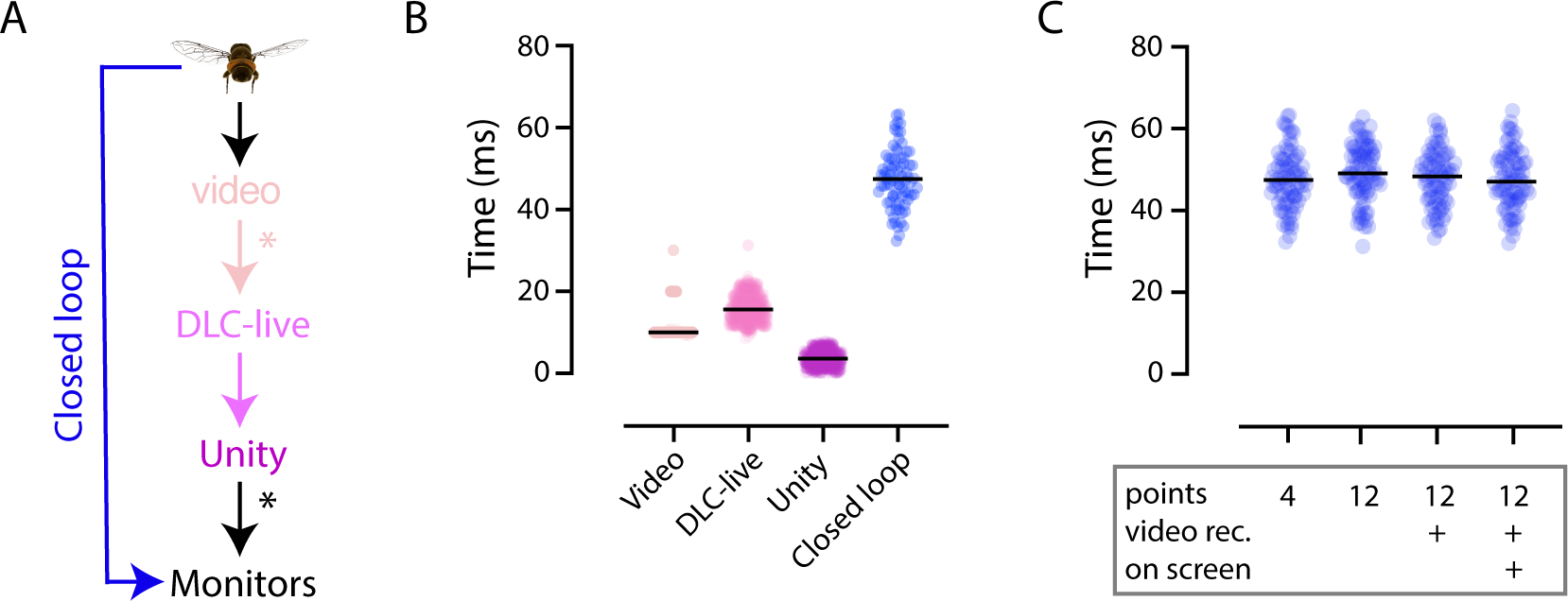
Closing the loop with low latencies. A) The tethered hoverfly is filmed from above, and each video frame sent to DLC-live. DLC-live performs markerless pose estimation and sends this information as packets to Unity. Unity calculates the WBA and the resulting scene update and sends this information to the monitors to close the loop. The stars indicate uncontrolled delays. B) The time taken for the steps described in panel A, using the same color coding. The video, DLC-live and Unity information was extracted from the DLC-Data file (see Fig. S1F). The time for closing the loop was measured with photodiodes. C) The time taken to close the loop under different conditions, where “points” refers to number of points tracked by DLC-live (e.g. colored dots, Fig. 1C), “video rec.” refers to whether the video was recorded or not, and “on screen” determines whether the video was visible or minimized during the experiment.

We next measured the time it took to close the loop using photodiodes and found this to be 47.6 ± 7.1 ms (mean ± std, blue, Fig. 8A, B), which is longer than the sum of the other three components (pink-scale, Fig. 8A, B). This suggests that the video camera does not immediately send each frame to DLC-live, that there is a delay between Unity drawing the frame and this being shown on the monitors, or a combination of the two (stars, Fig. 8A). Indeed, there are difficult to access latencies in conventional cameras and computer monitors (Stowers et al., 2014). Tracking more points, showing the DLC-live video on the screen, or simultaneously recording the video, did not make the process slower (Fig. 8C).

We measured the time it took to close the trackball loop using a video recording at 60 Hz. The peak in the cross-correlation between the pixel position of a vertical line on the screen and the rotation of the ball in the corresponding axis from FicTrac was a single frame. Given the 16.7 ms temporal resolution from recording at 60 Hz, this indicates the loop was effectively closed within one to two frames, i.e. within 16.7 – 33.3 ms.

## Discussion

We here present a flexible VR system developed by combining the gaming engine Unity with machine vision. We validated this with crabs walking on a trackball whose movements were recorded with FicTrac (Fig. 2, 4), and with tethered hoverflies where the wing movements were analyzed with DLC-live (Fig. 1, 7). In both cases, the actions of the tethered animal were used to control the movements of an avatar in a simulated environment (Fig. 3). We also developed a Unity user interface, CAVE, which allows easy control of yaw and gain settings (Fig. 5), experimental design (Fig. 6), and post-experiment analysis (Fig. S1). We show that the loop is closed within less than 50 ms (Fig. 8).

### Low delays for closing the loop

To make the VR immersive it is important to close the loop with low delays and to quantify this with photodiodes or similar. In humans, for example, cybersickness is induced when delays are longer than ca 60 ms, and the feeling of body ownership reduced when delays are above 100 ms (see e.g. Brunnström et al., 2020; Caserman et al., 2019). In VR, the total latency comes from three major components: tracking, rendering, and display (Stowers et al., 2014). Other VR systems, for example one that closed the loop based on centroid tracking of walking *Drosophila,* reported a 40 ms delay (Tadres and Louis, 2020), a Raspberry Pi Virtual Reality (PiVR) for optogenetic stimulation reported a 30 ms delay, and FlyVR has a latency of 40 – 80 ms (Stowers et al., 2014; Straw et al., 2011). Our measured delays were less than 50 ms (blue data, Fig. 8). While we found likely latencies in the camera and monitor transfer (stars Fig. 8A, and see Stowers et al., 2014), this could potentially be improved by using faster monitors or LED arrays, better CPU, graphics cards, or potentially different cameras. For example, the delay was lower in the trackball set-up using a FLIR camera.

Importantly, changing the number of points tracked by DLC-live, or how the videos were treated during the experiment, did not affect the latency for closing the loop (Fig. 8C). Furthermore, we show a mean delay of 47 ms (Fig. 8) allowed hoverflies to navigate towards flowers (Fig. 7) and with the even lower 17-33 ms delay, crabs to responded appropriately to other crabs (Fig. 4). However, for hoverflies, close to 50 ms may be too slow for faster behavioral interactions, including target pursuit (Fabian et al., 2018; Mischiati et al., 2015; Talley et al., 2023). As such, to allow pursuit behaviors or other fast conspecific interactions (Fig. 6D), this delay likely needs to decrease substantially.

One way to bypass this delay would be to implement a magnetic tether in the tethered flight configuration, allowing for unrestricted yaw turns (Duistermars and Frye, 2008), similar to our crab configuration (Fig. 2). This would allow the hoverfly to perform fast saccades, while the avatar’s translational motion is controlled by Unity. Alternatively, one could turn up the yaw gain (Fig. 5A-D) to allow for fast saccadic maneuvers. However, this is likely to induce flight instability, and might therefore not be practical.

### Usability

Our VR system using Unity with machine vision (DLC-live or FicTrac) is highly adaptable to different systems or species, as highlighted by our work using hoverfly tethered flight (Fig. 7) and crab trackball experiments (Fig. 4). Unity can be run by itself (Fig. 2-4), or with our user interface, CAVE, which provides a version that requires no coding to get started. CAVE allows the user to design visual experiments consisting of trials that can be put together into sequences (Fig. 3, 6, S1), and provides easy access to yaw and thrust gain settings (Fig. 5). Importantly, all data are stored in CSV files for offline analysis (Fig. S1F). This allows the user to combine detailed knowledge of the position of each OOI and the tethered animal’s avatar at each point in time (see e.g. Fig. 5I-K, 7A, B), which can be combined with offline analysis of the DLC movies, to quantify e.g. leg or head movements (Fig. 1C). Our GitHub repository includes not only user manuals, instruction videos and other documentation, but starting scenes, objects and OOIs, to get the novice user started.

While the supplied CAVE interface contains many useful tools, future improvements could e.g. include a repository of more naturalistic scenes from a range of habitats (Tolhurst et al., 1992). Furthermore, the monitors used here (Fig. 1A, 2A) are developed for human vision, and thus do not contain the correct color space for insects. In contrast, the Antarium (Kócsi et al., 2020) was developed to be optimized for ant vision. As such, in the future it could be beneficial to use a digital display that is more correct for insect vision, and which is updated with a higher refresh rate. We hope that by making our interface available on GitHub, other participants may contribute to such future development.

## CRediT author statement

**Yuri Ogawa:** methodology, validation, formal analysis, investigation, writing – review and editing, visualization, supervision; **Raymond Aoukar:** conceptualization, methodology, software, validation, formal analysis, investigation, writing – review and editing, visualization; **Richard Leibbrandt:** conceptualization, methodology, software, writing – review and editing; **Jake Manger**: methodology, software, validation, formal analysis, investigation, writing – review and editing, visualization; **Zahra Bagheri**: methodology, validation, formal analysis, investigation, supervision, writing – review and editing; **Luke Turnbull:** methodology, validation, investigation, writing – review and editing; **Chris Johnston:** methodology, software, writing – review and editing; **Pavan Kaushik:** conceptualization, methodology, software, writing – review and editing; **Jan Hemmi:** validation, resources, writing – review and editing, supervision, project administration, funding acquisition; **Karin Nordström:** conceptualization, validation, resources, writing – original draft, visualization, supervision, project administration, funding acquisition.

## Supporting information

Supp Movie 1. Crab in open loop and closed loop demonstration

Supp Movie 2. Validating yaw and thrust settings.

Supp Movie 3. Hoverfly in closed loop demonstration

## Acknowledgements

We thank Biomedical Engineering at SAHLN and the Botanic Gardens of Adelaide for their ongoing support. We thank Charlotte Goh for helping with running crab behavioral experiments and Juan Francisco Guarracino for helping with crab collection. This research was funded by the US Air Force Office of Scientific Research (AFOSR, FA9550-19-1-0294 and FA9550-23-1-0473), the Australian Research Council (ARC, DP180100491, FT180100289, DP200102642, DP210100740, and DP230100006) and the Flinders Foundation.

## Supp Figure

**Figure S1.**
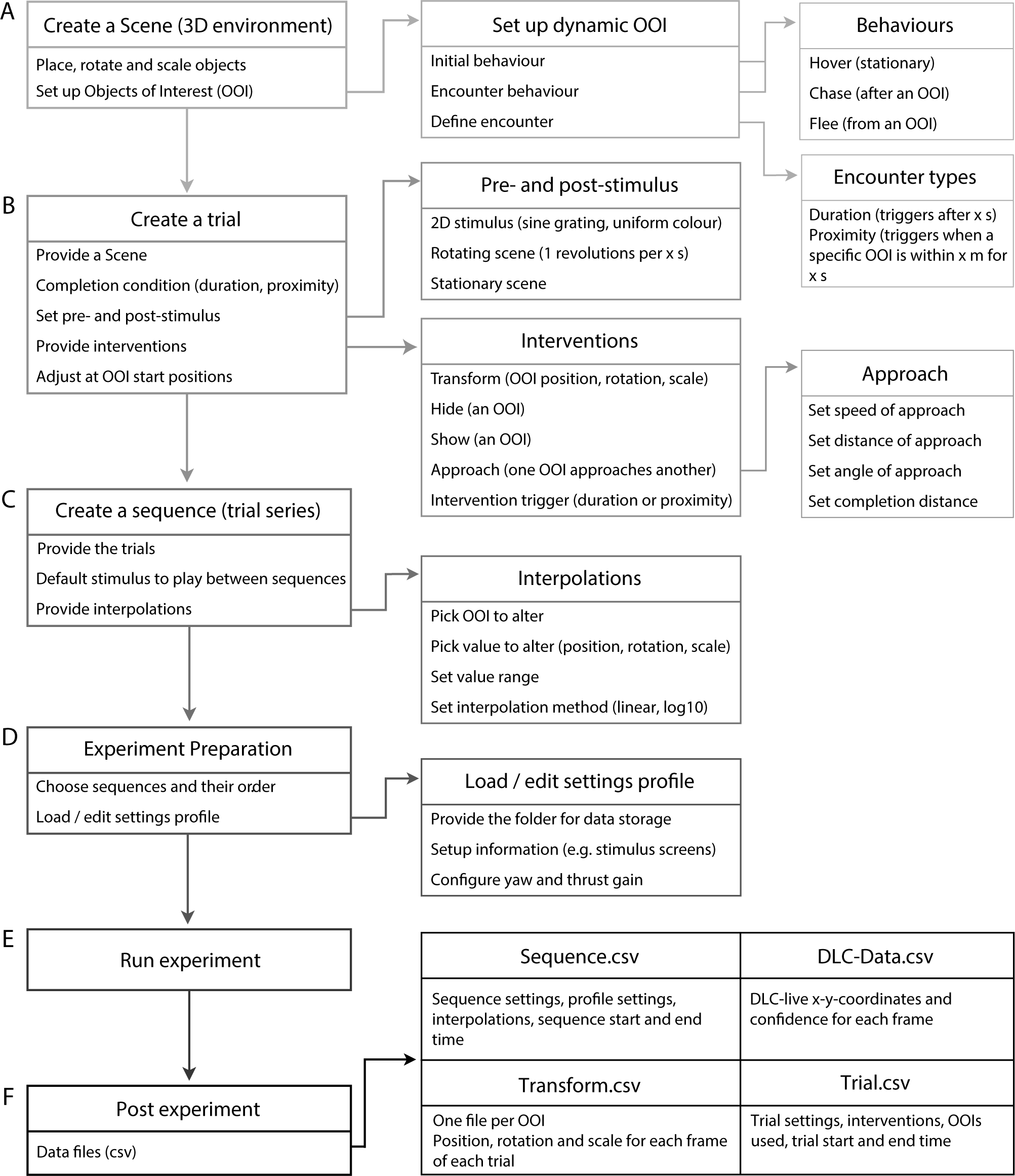
CAVE user story. A) The user chooses a scene and places objects within this. These can be defined as objects of interest (OOIs). If an OOI is dynamic, its behavior is defined. B) The user creates a trial, by using a scene, defining how a trial is completed, and what happens pre- and post-stimulus. At this stage, OOI interventions and their triggers are defined. C) A sequence consists of a series of trials, with a defined open loop stimulus in between each trial. Interpolations can be used to e.g. create OOIs of different sizes in each trial. D) The user choses what sequences to run, and loads the user specific settings, which include e.g. yaw and thrust settings. E) The experiment is run. F) Four data files are automatically created during the experiment. The sequence file contains all sequence information (as described in panel C), the DLC-Data files contain information about the DLC-live tracked points, the trial files contain all trial information (as described in panel B), and the transform file contains the x-z-location of the tethered animal’s avatar and each OOI for each frame of each trial.

## Supp Movies

**Supp Movie 1. Crab in open loop and closed loop demonstration.** The video shows a male *Gelasimus dampieri* responding distinctly to the visual stimuli. In open loop, it consistently avoids the virtual crab and briefly reacts to the virtual bird. In closed loop, the crab momentarily steps away from the virtual crab before evading the virtual bird.

**Supp Movie 2. Validating yaw and thrust settings.** This video was used to validate the yaw and thrust settings (Fig. 5I-K). The video was used as DeepLabCut-live input, which Unity then used to create the requisite scene updates.

**Supp Movie 3. Hoverfly in closed loop demonstration.** The video shows a female hoverfly in closed loop, navigating a scene with two dandelions, as in Fig. 7. The main part of the video shows the unwrapped view across the three screens. The top left inset shows the video and the points tracked by DLC-live in real time. The top right inset shows the virtual scene as seen from above, with the location of the flowers as yellow circles, and their proximity radii as dashed circles.

